# Molecular Specializations Underlying Phenotypic Differences in Inner Ear Hair Cells of Zebrafish and Mice

**DOI:** 10.1101/2024.05.24.595729

**Authors:** Kimberlee P. Giffen, Huizhan Liu, Kacey L. Yamane, Yi Li, Lei Chen, Ken L. Kramer, Marisa Zallocchi, David Z.Z. He

**Affiliations:** Department of Neuroscience and Regenerative Medicine, Medical College of Georgia at Augusta University, Augusta, GA, USA; Augusta University/University of Georgia Medical Partnership, Athens, GA, USA; Department of Biomedical Sciences, Creighton University School of Medicine, Omaha, NE, USA; Department of Otorhinolaryngology, Beijing Tongren Hospital, Beijing Capital Medical University, Beijing, China; Department of Cell and Developmental Biology, Vanderbilt University, Nashville, TN, USA

**Keywords:** hair cells, orthologs, RNA-seq, transcriptome, prestin, zebrafish, mouse, hearing

## Abstract

Hair cells (HCs) are the sensory receptors of the auditory and vestibular systems in the inner ears of vertebrates that selectively transduce mechanical stimuli into electrical activity. Although all HCs have the hallmark stereocilia bundle for mechanotransduction, HCs in non-mammals and mammals differ in their molecular specialization in the apical, basolateral and synaptic membranes. HCs of non-mammals, such as zebrafish (zHCs), are electrically tuned to specific frequencies and possess an active process in the stereocilia bundle to amplify sound signals. Mammalian cochlear HCs, in contrast, are not electrically tuned and achieve amplification by somatic motility of outer HCs (OHCs). To understand the genetic mechanisms underlying differences among adult zebrafish and mammalian cochlear HCs, we compared their RNA-seq-characterized transcriptomes, focusing on protein-coding orthologous genes related to HC specialization. There was considerable shared expression of gene orthologs among the HCs, including those genes associated with mechanotransduction, ion transport/channels, and synaptic signaling. For example, both zebrafish and mouse HCs express *Tmc1, Lhfpl5, Tmie, Cib2, Cacna1d, Cacnb2, Otof, Pclo* and *Slc17a8*. However, there were some notable differences in expression among zHCs, OHCs, and inner HCs (IHCs), which likely underlie the distinctive physiological properties of each cell type. *Tmc2 and Cib3* were not detected in adult mouse HCs but *tmc2a* and *b* and *cib3* were highly expressed in zHCs. Mouse HCs express *Kcna10*, *Kcnj13*, *Kcnj16*, and *Kcnq4*, which were not detected in zHCs. *Chrna9* and *Chrna10* were expressed in mouse HCs. In contrast, *chrna10* was not detected in zHCs. OHCs highly express *Slc26a5* which encodes the motor protein prestin that contributes to OHC electromotility. However, zHCs have only weak expression of *slc26a5*, and subsequently showed no voltage dependent electromotility when measured. Notably, the zHCs expressed more paralogous genes including those associated with HC-specific functions and transcriptional activity, though it is unknown whether they have functions similar to their mammalian counterparts. There was overlap in the expressed genes associated with a known hearing phenotype. Our analyses unveil substantial differences in gene expression patterns that may explain phenotypic specialization of zebrafish and mouse HCs. This dataset also includes several protein-coding genes to further the functional characterization of HCs and study of HC evolution from non-mammals to mammals.

## Introduction

Hair cells (HCs) are the sensory receptors of the auditory and vestibular systems in the inner ear of all vertebrates. HCs transduce mechanical stimuli, i.e., movement in their environment, into electrical activity (Hudspeth and Corey, 1977). The site of such mechanoelectrical transduction is the stereocilia bundle, the hallmark of all HCs. In addition to the hair bundle on the apical membrane, the basolateral and synaptic membranes of HCs have structural and functional specializations which are responsible for electrical activities and synaptic transmission.

Even though mechanotransduction in the hair bundle is a shared feature of all HCs, there are profound differences in morphological and functional specializations between HCs of non-mammals and mammals. The HCs of non-mammalian species, such as fish and birds, enhance their auditory sensitivity and frequency selectivity via an electrical resonance shaped by voltage-gated ion channel activity and an active, force-generation process in the stereocilia bundle (Fettiplace and Fuchs, 1999; Salvi et al., 2015; Nicolson, 2017; Fettiplace, 2020). Mammals have evolved two morphologically and functionally distinct HC types, the inner HCs (IHCs) and the outer HCs (OHCs). In contrast to non-mammals, the mammalian cochlear HCs are not electrically tuned, but rather accomplish the high sensitivity and frequency selectivity of hearing by using the mechanically tuned basilar membrane and prestin-based somatic motility of OHCs to amplify mechanical signals in the cochlea (Liberman et al., 2002; Dallos et al., 2008; Hudspeth, 2014). The morphological and physiological differences among vertebrate HCs are still being explored. Comparison of the transcriptomic signatures underlying phenotypic specializations of non-mammalian and mammalian HCs can further our understanding of the molecular mechanisms underlying these differences.

Previously, we isolated pure populations of adult zebrafish inner ear HCs and adult mouse IHCs and OHCs for RNA-seq-based transcriptomic analyses (Barta et al., 2018; Li et al., 2018). To analyze the molecular differences among these HC populations, the transcriptomic datasets were refined to include only protein-coding gene orthologs. Our previous study identified 17,498 protein-coding gene orthologs in zebrafish which correspond to 13,557 orthologous mouse genes (Giffen et al., 2019). This study aimed to characterize the expressed genes either common or unique to the HCs of each species, focusing on the genes known to encode HC specializations in the apical, basolateral, and synaptic membranes, since these genes should largely be responsible for phenotypical differences between zebrafish and mammalian HCs. Additionally, we compared expression of genes with a known association to hearing loss. Genes encoding transcription factors were also examined to elucidate pathways regulating expression of these protein-coding gene orthologs. Our transcriptome analyses are expected to serve as a highly valuable resource, not only for unraveling the molecular mechanisms of the unique biological properties of zebrafish and mouse HCs, but also to further our understanding of HC evolution from lower vertebrates to mammals.

## Materials and Methods

### Zebrafish and Mouse protein-coding gene orthologs

The built-in Ensembl Biomart package was used to access the annotated Ensembl Genes database (release 91) (Cunningham et al., 2022) to generate a list of zebrafish and mouse protein-coding gene orthologs. The Biomart classification and quality of orthologs incorporates a gene order conservation, both up and downstream of the gene of interest, whole genome alignment, and conservation of nucleotide sequences to predict high-confidence orthology thresholds (Zerbino et al., 2018). The annotated gene lists for this study were based on the Zebrafish (*Danio rerio*) GRCz10 and mouse (*Mus musculus)* GRCm38 genome builds. In the zebrafish-to-mouse comparison, the zebrafish genes were filtered based on protein-coding function and homology with mouse genes, producing a total of 17,498 orthologs. Similarly, a mouse-to-zebrafish comparison identified a total of 13,526 protein-coding gene orthologs. The final mouse gene list included fewer orthologs because redundant genes corresponding to multiple zebrafish paralogs were removed from the gene list for analysis.

### RNA-seq datasets

Previously published RNA-seq datasets were used in the comparative analysis of expressed protein-coding genes. Adult *Tg(pou4f3:GAP43-GFP)s273t* zebrafish inner ear HCs and non-sensory supporting cells (SCs) were collected using a unique suction pipette technique and isolated RNA was sequenced for transcriptomic analysis (He et al., 2000; Liu et al., 2014; Barta et al., 2018) (NCBI SRP113243). Similarly, isolated populations of IHCs and OHCs from one-month-old CBA/J mouse cochleae were collected and utilized for RNA-seq analysis (Li et al., 2018) (NCBI SRP133880).

The quantified cell-type specific gene expression data from these previous studies and the protein-coding gene ortholog lists were merged into a single dataset, which is provided as a searchable Excel table (Supp Data Sheet 1). The arbitrary expression value of 0.1 RPKM (Reads per kilobase per million mapped reads) (FDR *p*-value ≤ 0.10) was set as the minimum expression cutoff for both species. The expression values presented herein are not quantitatively equivalent, but rather are indicative of relative expression in mouse and zebrafish inner ear cell populations and thus should not be compared as absolute values. No custom code was used in this analysis.

### Gene enrichment analysis

To further characterize the functions of groups of commonly or uniquely expressed gene orthologs, gene enrichment analysis was conducted using ShinyGO v 0.80 (Ge et al., 2020). This web-based program utilizes the annotated Gene Ontology (GO) and other databases to provide enrichment analysis results. The *p*-value was set at < 0.05.

### Validation of gene expression

RT-PCR: A number of genes were selected for verification by RT-PCR based on shared or differential expression in zebrafish and mouse HCs. The oligonucleotide primers were designed with A plasmid Editor (ApE) software to find unique and appropriate sequences (Davis and Jorgensen, 2022). Cell populations were collected using previously published protocols (Liu et al., 2014; Barta et al., 2018). Total RNA was isolated using the Qiagen miRNeasy kit and quantified using a nanodrop spectrophotometer. The cDNA libraries were prepared from isolated RNA with the iSCRIPT reverse transcription supermix (Bio-Rad). RT-PCR reactions were prepared as 20 μl reactions using 2x Master mix (Bio-Rad). All primers (Appendix 1) were acquired from Integrated DNA Technologies (Coralville, Iowa). Mouse stria cells and zebrafish non-sensory SCs were included as controls.

smFISH: To examine mRNA expression of select genes in the adult zebrafish and mouse inner ear epithelia, we used the RNAscope-based single molecule fluorescent in situ hybridization (smFISH) assay from Advanced Cell Diagnostics (ACD). Samples were pretreated according to the ACD protocol for formalin-fixed paraffin-embedded tissue. The pretreatment target retrieval and protease steps were slightly modified to prevent tissue damage. Tissues were covered with freshly prepared 0.5% pepsin + 5 mM HCl in deionized water and incubated in the humidifying chamber at 37°C for 10 minutes. The remaining protocol was conducted according to the RNAscope 2.5 HD Detection Reagent RED user manual and published protocols (Salehi et al., 2018). Following the final wash, samples were incubated with 4′,6-diamidino-2-phenylindole (DAPI) to label nuclei. The smFISH probes were detected by red channel fluorescence (imaging described below), with no probe treatment included as a negative control.

Immunofluorescence: Mice (WT C57BL/6J) and zebrafish *Tg(Pou4f3:,GFP*) were euthanized according to IACUC protocols. Mouse otic capsules and zebrafish inner ear tissues were dissected and placed in 4% paraformaldehyde (PFA) in 1X phosphate buffered saline (PBS) for 24 hours at 4 °C. Following dissection for whole mount preparations, samples were washed in 1X PBS, then permeabilized and blocked in 1X PBS with 0.2% Triton X-100 (PBS-T) and 5% normal goat serum (NGS), followed by incubation with SLC7A14 primary antibody (Sigma:HPA045929; lot:R43519; 1:400) overnight at 4 °C. Samples were rinsed in 1X PBS three times and then incubated with Alexa-Fluor (AF) conjugated secondary antibodies (Invitrogen) for one hour at room temperature followed by subsequent washes. Additional staining with AF488-Phalloidin (anti-F-Actin) was used to label stereocilia. Finally, samples were rinsed three times in 1X PBS and mounted with SlowFade (Invitrogen), then imaged as below.

Confocal microscopy: Visualization of smFISH and immunofluorescence was conducted using Zeiss laser scanning confocal microscopes LSM700, 710 or 880. Sections were imaged at 1.0 zoom at 40x or 63x magnification using Zen Black acquisition software. Additionally, samples for colocalization analyses, included cultured cells, were imaged on a Nikon Ti-E with a Yokogawa Spinning Disc Confocal with a Flash 4.0 Hamamatsu Monochrome camera.

### Simultaneous Recording of Motility and Nonlinear Capacitance under Voltage-Clamp Condition

Solitary HCs from zebrafish and mice were placed in the extracellular solution contained (mM): 120 NaCl, 20 TEA-Cl, 2 CoCl_2_, 2 MgCl_2_, 10 HEPES, and 5 glucose. Whole-cell voltage-clamp recordings were performed on an Olympus inverted microscope (IX71) with DIC. The patch pipette with the headstage of an Axopatch 200B amplifier (Axon Instruments) was held by a Narashige 3D micromanipulator (MHW-3). Whole cell voltage-clamp tight-seal recordings were established. The patch electrodes were pulled from 1.5 mm glass capillaries (WPI) using a Flaming/Brown Micropipette Puller (Sutter Instrument Company, Model P-97). Recording pipettes had open tip resistances of 4-5 MΩ and were filled with an internal solution that consisted of (mM): 140 CsCl, 2 MgCl_2_, 10 EGTA, and 10 HEPES. The osmolarity and pH for both intracellular and extracellular solutions were adjusted to 300 mOsm/l and 7.3. The access resistance typically ranges from 8 to 14 MΩ after the whole-cell recording configuration was established. Voltage error due to the uncompensated series resistance was compensated off-line. All experiments were performed at room temperature (22±2^0^C)

Motility and NLC were acquired simultaneously as described elsewhere (Wang et al., 2010). The AC technique was used to obtain motility-related gating charge movement and the corresponding nonlinear membrane capacitance (Santos-Sacchi et al., 1998). In brief, it utilized a continuous high-resolution (2.56 ms sampling) two-sine voltage stimulus protocol (10 mV for both 390.6 and 781.2 Hz), with subsequent fast Fourier transform-based admittance analysis. These high frequency sinusoids were superimposed on staircase voltage (in 4 mV steps) stimuli. The evoked capacitive currents were filtered at 5 kHz and digitized at 100 kHz using jClamp software (SciSoft Company), running on an IBM-compatible computer and a 16-bit A/D converter (Digidata 1322, Molecular Devices). Motility was measured and calibrated by a photodiode-based measurement system (Jia and He, 2005) mounted on the Olympus inverted microscope. The magnified image of the edge of the cell was projected onto a photodiode through a rectangular slit. Somatic length changes, evoked by voltage stimuli, modulated the light influx to the photodiode. The photocurrent response was calibrated to displacement units by moving the slit a fixed distance (0.5 µm) with the image of the cell in front of the photodiode. After amplification, the photocurrent signal was low-pass filtered by an anti-aliasing filter before being digitized by a 16-bit A/D board (Digidata 1322, Molecular Devices). The photodiode system had a cutoff (3 dB) frequency of 1100 Hz. The motile responses were filtered at 1100 Hz and digitized at 100 kHz. The voltage dependence of the cell length change also fits with the two-state Boltzmann function and can be described as:

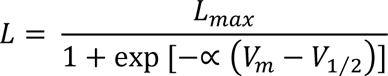

where *L*_max_ is the maximal length change, *V*_1/2_ is the voltage at which *L* = 0.5*L*_max_, *α* is the slope factor indicating the voltage sensitivity of the length change. Slope function was obtained as the derivative of the Boltzmann function. The nonlinear capacitance can be described as the first derivative of a two-state Boltzmann function relating nonlinear charge movement to voltage (Ashmore, 1989; Santos-Sacchi, Joseph, 1991). The capacitance function is described as:

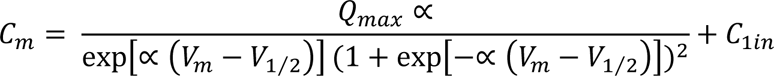

## Results

### Orthologous relationships of zebrafish and mouse protein-coding genes

A direct comparison of zebrafish and mouse protein-coding genes was conducted using Ensembl Biomart, based on the zebrafish GRCz10 and mouse GRCm38 gene assemblies (Cunningham et al., 2022). The analysis revealed a high number of orthologous genes shared among the two species (Table 1). Among the total zebrafish protein-coding genes, almost 70 percent had at least one mouse ortholog. The total number of orthologous zebrafish genes is greater as a result of a known ancient genome duplication event, so there were redundant mouse genes mapped to paralogous zebrafish genes (i.e., many-to-one, many-to-many) (Koonin, 2005). The one-to-one zebrafish-to-mouse orthologs accounted for 39 percent of the zebrafish protein-coding genes. Furthermore, 3,248 of these genes were identified as high-confidence orthologs, with a gene order conservation score of ≥ 75 and/or whole genome alignment score of ≥ 75 and a percent identity score of ≥ 50.

**Table 1:** Classification of orthologous protein-coding genes.

| **Relationship** | **Zebrafish** | **Mouse** |
| --- | --- | --- |
| One-to-one | 9,864 |  |
| One-to-many | 6,723 | 3,425 |
| Many-to-many | 911 | 237 |
| Ortholog total | 17,498 | 13,526 |
| Unique | 7,600 | 8,532 |
| Protein- coding gene total | 25,098 | 22,058 |

### Commonly and uniquely expressed protein-coding gene orthologs among zebrafish and mouse inner ear HC populations

An initial comparison of the number of expressed protein-coding gene orthologs in the zHCs showed that 10,995 of the 17,498 orthologs were detected above the cutoff expression value of 0.10 RPKM (FDR *p*-value ≤ 0.10) (Figure 1). Of the 13,526 mouse gene orthologs, 8,547 were expressed in IHCs, while 8,863 were expressed in OHCs (Figure 1). Further comparison showed that a total of 6,659 genes were commonly expressed among all sensory HCs, with an additional 932 zebrafish gene paralogs. There were 149 genes (+25 paralogs in zebrafish) commonly expressed in zHCs and IHCs, while there were 322 genes (+77 paralogs in zebrafish) expressed in zHCs and OHCs. The mouse IHCs and OHCs shared a greater number of commonly expressed genes. The zHCs had the highest number of uniquely expressed genes (2,831), while the IHCs and OHCs expressed 145 and 288 unique genes, respectively.

**Figure 1:**
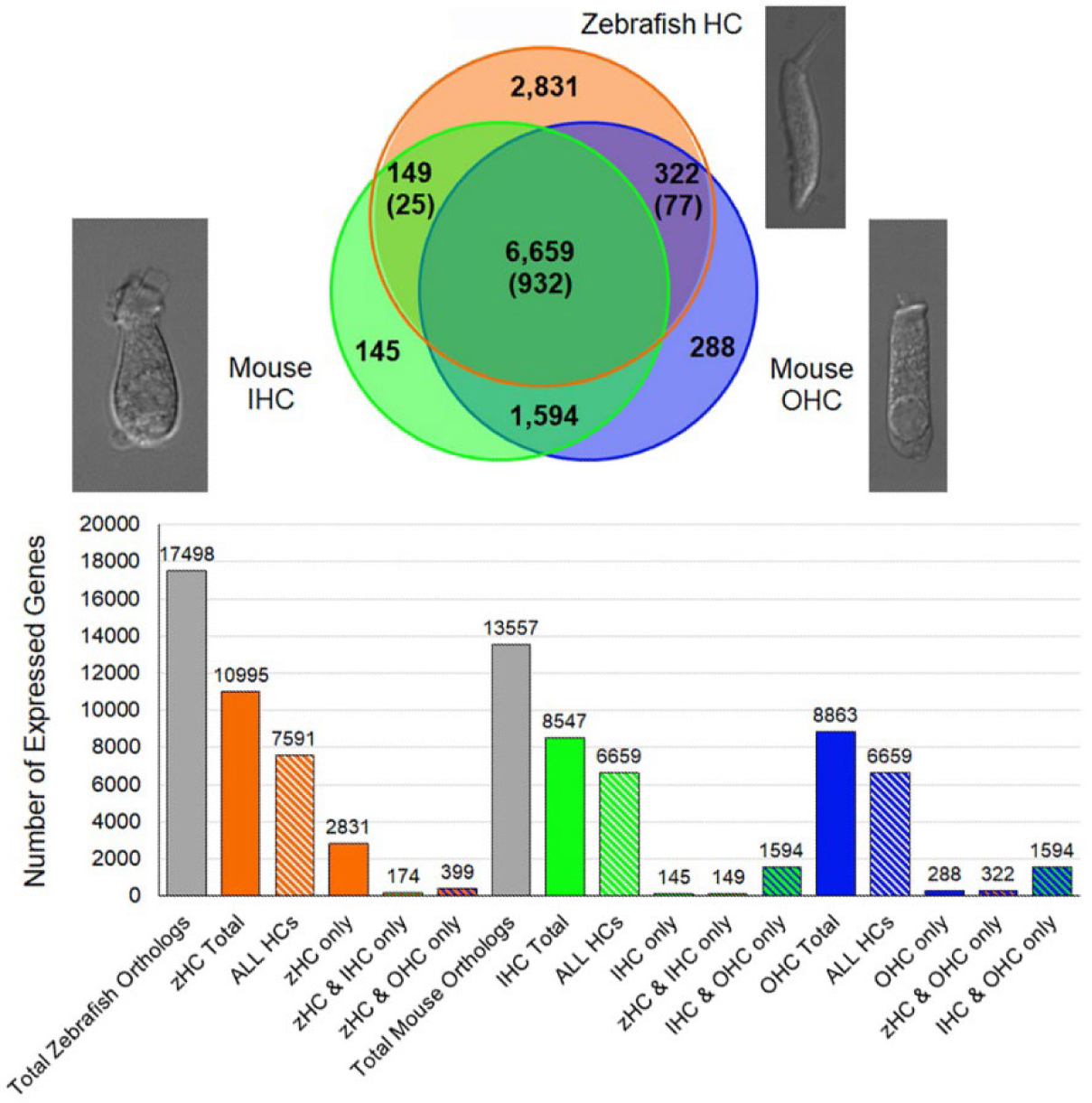
Protein-coding gene orthologs expressed in zebrafish and mouse HCs. Commonly and uniquely expressed protein-coding gene orthologs among zHC, and mouse IHC and OHC populations are shown in the Venn diagram. The number in parentheses indicates additional paralogous genes expressed only in zHCs. The bar graph shows the number of uniquely and commonly expressed gene orthologs among the HC populations.

### Enriched protein-coding gene orthologs expressed in zebrafish inner ear HCs compared to non-sensory zSCs

We analyzed differentially and uniquely expressed genes in zHCs with reference to non-sensory zSCs from the zebrafish inner ear since many of these genes may underlie specialization of sensory HCs. The top 200 gene orthologs uniquely expressed in zHC (zSCs ≤ 0.10 RPKM, FDR *p*-value ≤ 0.10) are shown in figure 2A. Several of these genes are known to contribute to essential HC function in zebrafish including *chrna9a/b*, *lhflp5a*, *myo7aa*, *strc*, and *tmie* (Erickson and Nicolson, 2015; Nicolson, 2017). The top 200 upregulated, differentially expressed genes in zHCs (> 0.1 RPKM in zHCs and zSCs; Log2 fold change ≥ 1.0; FDR *p*-value ≤ 0.10) are shown in figure 2B, with the fold-change in expression relative to the zSCs, shown to the right of each panel. Consistent with other studies, several of the top differentially expressed genes are pan-HC genes including *otofb, slc17a8,* and *ush1c* (Erickson and Nicolson, 2015; Erickson et al., 2019; Shi et al., 2023).

**Figure 2:**
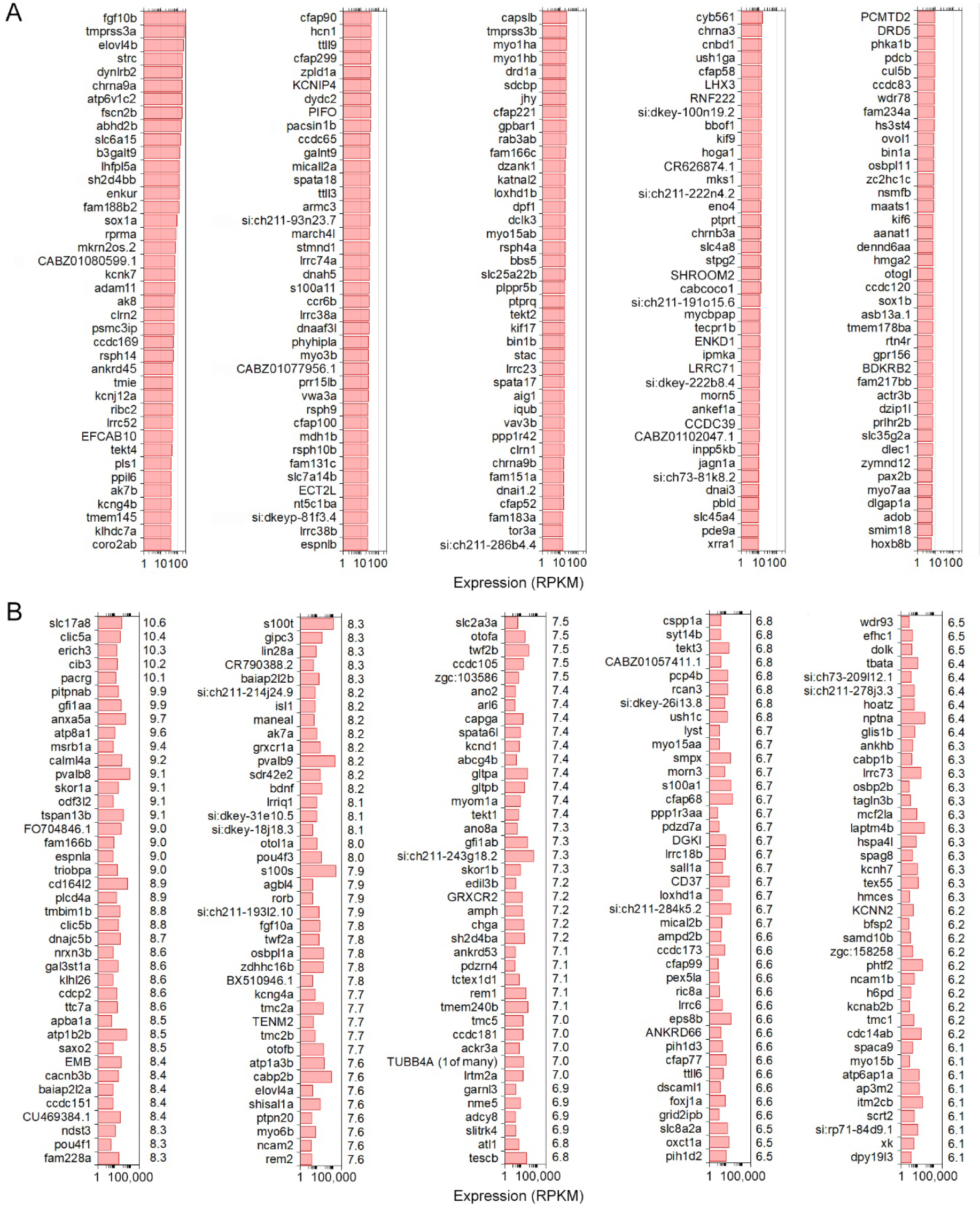
Uniquely and differentially expressed genes in zHCs. **A**: The top 200 uniquely expressed gene orthologs in zHCs (in red) (zSCs ≤ 0.10 RPKM, FDR *p*-value ≤ 0.10). **B**: The top 200 upregulated, differentially expressed genes in zHCs (in red) with comparison to zSCs (in grey) (> 0.1 RPKM in zHCs and zSCs; Log2 fold change ≥ 1.0; FDR *p*-value ≤ 0.10). Numbers on the right side of each panel are Log2-fold change in expression, relative to the zSCs.

### Gene ortholog expression among zHCs and mouse IHCs

Next, we examined similarities among the transcriptomic signatures of the primary auditory receptor cells in the mouse cochlea, IHCs, and their homolog in the zebrafish inner ear sensory epithelia. The top 200 expressed protein-coding orthologs in zHCs and IHCs are shown in Figures 3A and 3B, respectively. Among these top expressed genes, there were 47 genes commonly expressed in all HCs, 27 of which were one-to-one gene orthologs and 25 were ranked as high-confidence orthologs. Many of the highly expressed genes are associated with metabolic processes, including mitochondrial genes. Genes commonly expressed among the zHCs and IHCs including *anxa5a* (*Anxa5*), *pvalb8* and pvalb9 (*Ocm*) and *otofb* (*Otof*).

**Figure 3:**
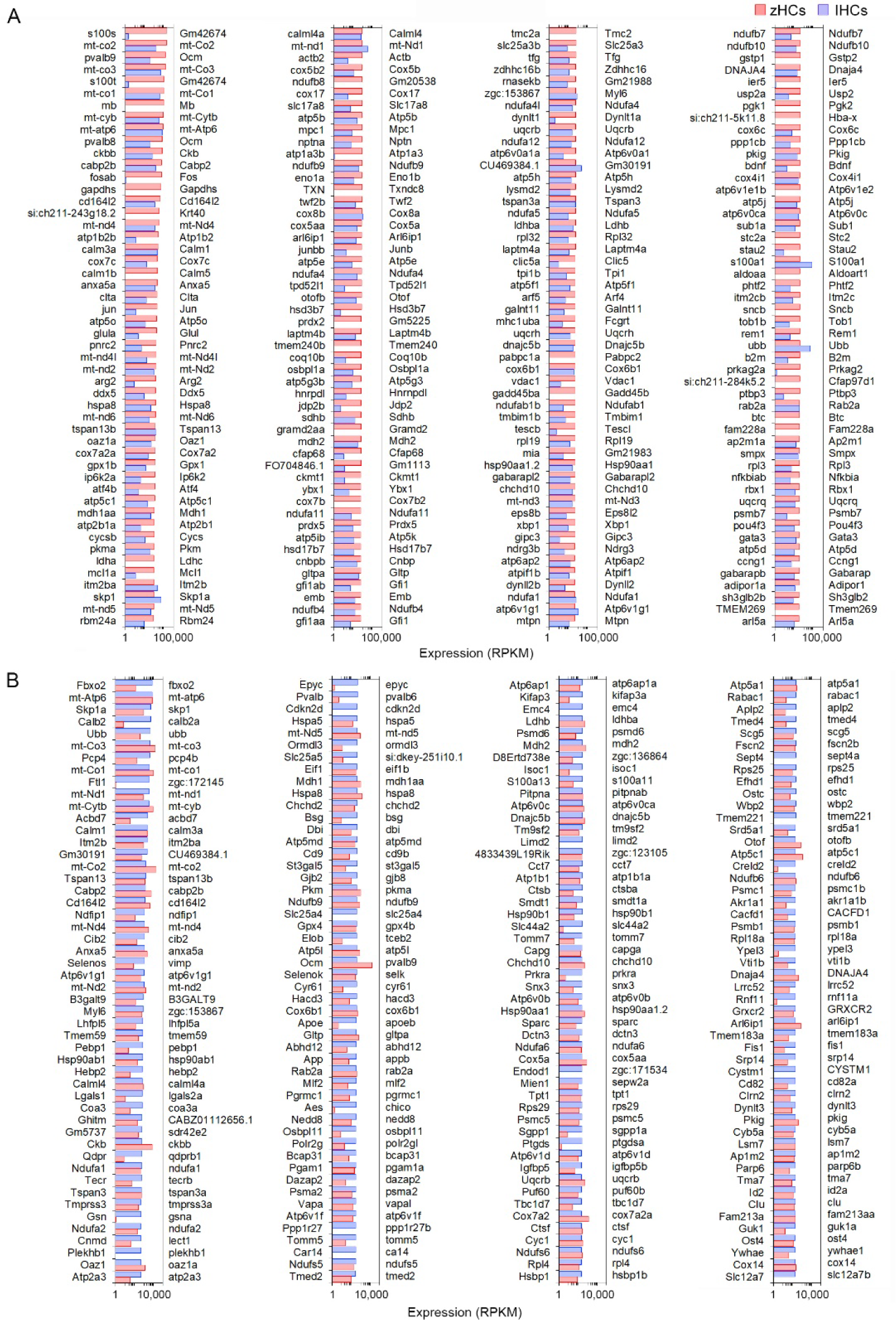
Comparison of the top 200 protein-coding gene orthologs expressed in zHCs and mouse IHCs. **A**: Top 200 expressed gene orthologs in zHCs (≥ 0.10 RPKM, FDR p-value < 0.10). Expression of gene orthologs in mouse IHCs (gene symbols listed on the right side of each panel) presented for comparison. **B**: Top 200 gene orthologs expressed in IHCs (≥ 0.10 RPKM, FDR p-value < 0.10), with expression of the gene orthologs in zHCs presented for comparison. Note that direct comparison of normalized RPKM values is not reliable as the datasets for each species were analyzed independently. Legend included in top right of figure.

### Expression of genes encoding functional HC proteins

Vertebrate HCs have specializations in the apical, basolateral, and synaptic membranes that are responsible for mechanotransduction, electrical and mechanical activities, and synaptic transmission. To examine expression of gene orthologs encoding proteins associated with the stereocilia bundle, we first generated a gene list based on two studies that examined proteins present in stereocilia bundles of mouse vestibular and chicken HCs using mass spectrometry (Shin et al., 2013; Krey and Barr-Gillespie, 2018). Fig. 4A shows expression levels of 113 gene orthologs, 45 of which are one-to-one orthologs, that encode stereocilia-associated proteins. As shown, 86 of the genes expressed in mammalian HCs are also expressed in zHCs, specifically those encoding the cytoskeletal structure and associated proteins. One-to-one gene orthologs highly expressed among all HC types included *arpc2, atp5b, capzb, cdc42, cfl1, fbxo2, gdi2, grxcr2, twf2b* (*Twf2*), and *ush1c*. Other distinctive patterns in gene expression among the HC populations were also identified. For example, *ldhba* (*Ldhb*) and *pvalb9* (*Ocm*) were highly expressed in all HCs, though notably, zHC and OHCs had higher expression than IHCs; *ldhba* (*Ldhb*) has been implicated in age-related hearing loss (Tian et al., 2020). *Actn4* and *Vill* were expressed only in IHCs and OHCs, not zHCs. Expression of both *capza1a and 1b* was observed in zHCs, while *capzb* (*Capzb*) was highly expressed in all HCs. Finally, there was notable redundancy in expression of paralogous genes in zHCs including *calm1a* and *b, dynll2a* and *b*, *pcdh15a* and *b*, etc.

**Figure 4:**
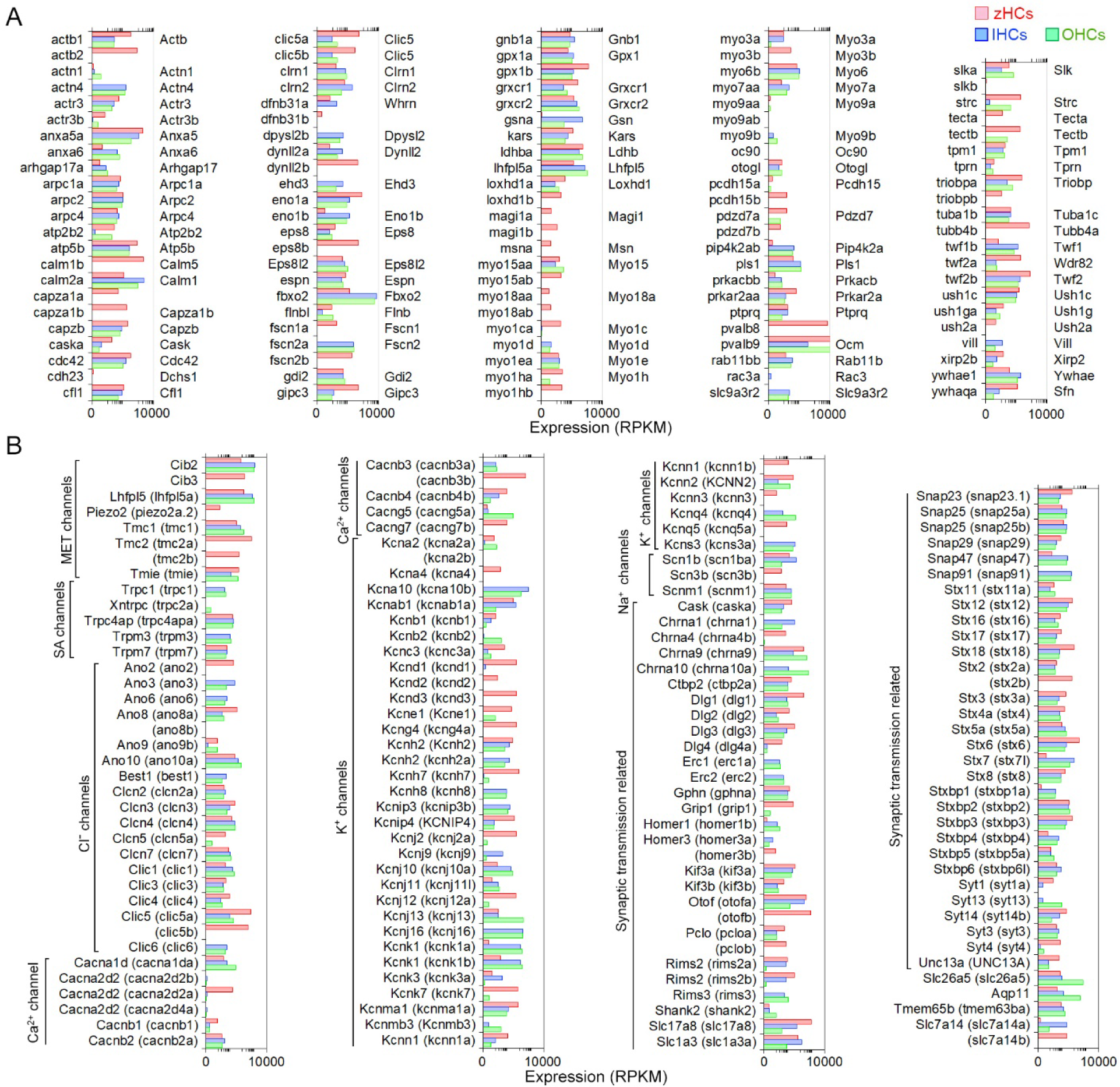
Genes associated with HC specialization and function. **A**: Expression of gene orthologs associated with the HC stereocilia bundle. The gene list was generated from the proteins detected from the stereocilia bundle preparations of chicken HCs and mouse vestibular HCs, then reduced to 113 protein-coding gene orthologs. The zHC genes are listed to the left of the panel with mouse orthologs listed to the right of the panel. Gene expression in zHCs, IHCs, and OHCs is shown, with a color-coded legend included on top of the figure. **B**: Comparison of gene orthologs encoding proteins associated with HC function including mechanotransduction, ion transport, synaptic transmission, motor function, and other functions. Zebrafish gene name in parentheses. Legend included in top right of figure.

Next, we further examined other HC function-associated genes, including those encoding components of the mechanotransduction apparatus. All HCs contain mechanotransduction channels in the stereocilia, including TMC1 (transmembrane channel-like 1) and/or TMC2 (Kawashima et al., 2011; Kim et al., 2013; Pan et al., 2013). LHFPL5, TMIE, and PIEZO2 are also necessary for mechanotransduction (Kalay et al., 2006; Xiong et al., 2012; Zhao et al., 2014; Wu et al., 2017). As shown in Fig. 4B, *Tmc1* (*tmc1*) is highly expressed in mouse HCs and zHCs. *Tmc2* is not detected in adult mouse HCs; however, both *tmc2a* and *tmc2b* are highly expressed in zHCs with expression values greater than *tmc1*. *Lhfpl5* (*lhfpl5a*) and *Tmie* (*tmie*) are expressed in mouse and zebrafish HCs, respectively.

HCs also contain various types of ion channels in the basolateral and synaptic membranes that are responsible for establishing and maintaining membrane potential, shaping receptor potentials, facilitating neurotransmitter release, and regulating cell volume (Kros, 1996; Mittal et al., 2017). Fig. 4B includes expression of gene orthologs that encode various types of Cl^−^, Ca^2+^, Na^+^, and K^+^ channels. Differential expression of several of these ion channel genes in IHCs and OHCs was previously observed in our RNA-seq analysis (Liu et al., 2014). There are some distinct differences in the expression of ion channel-related genes between zebrafish and mouse HCs. Zebrafish HCs have robust expression of *cacnb3b* (encoding voltage-dependent L-type calcium channel subunit beta-3), while mouse HCs have weak expression of *Cacnb3*. Conversely, robust expression of *Cacna1d* (encoding voltage-dependent calcium channel, L type, alpha 1 subunit) and *Cacng5* (encoding voltage-dependent calcium channel, gamma subunit 5 or transmembrane AMPAR regulatory protein gamma-5) are detected in mouse HCs, especially in OHCs. Relatively high-level expression of *kcnd1*, *kcnd3*, *kcng4a, kcnh7, kcnj12a, kcnk7*, *kcnma1a*, and *kcnq5a* is detected in zHCs. In contrast, robust expression of *Kcnj13* and *Kcnq4* is detected in OHCs, IHCs have relatively high expression of *Kcnh2, Kcnk1,* and *Kcns3*, and both express *Kcnj10, Kcnj16,* and *Kcnk1*. Also, there was similar expression of *kcnab1a* (*Kcnab1*) in zHCs and IHCs, and *kcnn2* (*Kcnn2*) in zHCs and OHCs.

Finally, we examined expression of genes that encode proteins associated with pre-synaptic and post-synaptic mechanisms including pre-synaptic neurotransmitter vesicle transport and release, as well as post-synaptic nicotinic cholinergic receptors in zebrafish and mouse HCs. As shown in Fig. 4B, the expression of genes encoding synaptic proteins is similar in zebrafish and mouse HCs, while differential expression of genes may underlie distinct synaptic morphology and function of HC populations. For example, *Otof* (*otofa/b*) (Roux et al., 2006)*, Slc17a8* (*slc17a8*) (Ruel et al., 2008) and *Slc1a3* (*slc1a3a)* (Liu et al., 2014; Barta et al., 2018) are more highly expressed in IHCs and zHCs. Interestingly, while both *Chrna9* (Elgoyhen et al., 1994) and *Chrna10* (Vetter et al., 2007) are expressed in mouse HCs, especially OHCs, zHCs only express *chrna9*.

### Expression of gene orthologs associated with a hearing loss phenotype

A subset of genes associated with hearing loss phenotypes was compiled based on a list in our previous publication (Liu et al., 2014). Expression of 115 protein-coding mouse genes and corresponding zebrafish gene orthologs, 34 of which are classified as high confidence orthologs, is shown in Figure 5. There were several gene orthologs that were highly expressed in zHCs, IHCs, and OHCs including *Cib2*, *Gjb2* (*cx30.3*), *Grxcr2*, *Lhfpl5* (*lhfpl5a*), *Smpx*, and *Wpb2*. The gene ortholog *Strc*, encoding the stereocilin protein which forms horizontal connections in OHC stereocilia (Cartagena-Rivera et al., 2019), was highly expressed in zHCs and OHCs. The orthologs *Slc17a8*, *Slc7a14* (*slc7a14b*), and *Otof* (*otofb*) were highly expressed in zHC and IHC.

**Figure 5.**
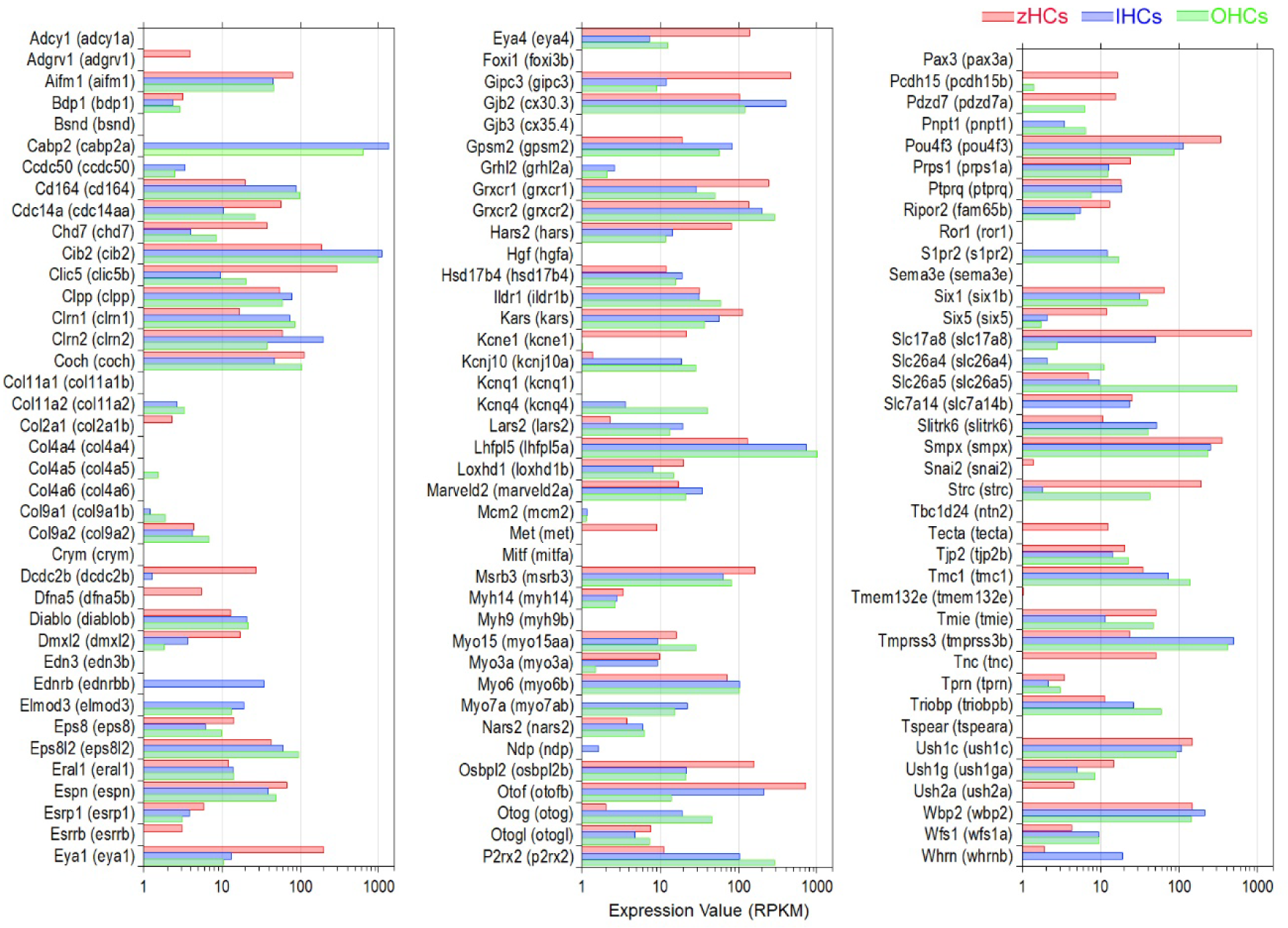
Expression of protein-coding gene orthologs identified in hereditary deafness. This gene list was based on a list compiled in our previous publication (Liu et al., 2014). Zebrafish gene name in parentheses. Legend included in top right of figure.

### Common expression of gene orthologs encoding transcriptional regulators

To identify possible regulatory mechanisms that maintain mature vertebrate HC phenotypes, common expression of genes with transcription-associated functions was examined. An inclusive list of over 2,300 “transcription factor” (TF) gene orthologs, curated in a previous publication (Giffen et al., 2019) included those with functions such as regulatory region DNA binding, chromatin-mediated transcriptional regulation, nuclear binding activity, and similar functions. First, to understand the conserved transcriptomic signatures of vertebrate HCs, the high-confidence gene orthologs expressed in all HCs were examined as these TF genes are more likely to have conserved functions. A total of 400 high-confidence TF gene orthologs were expressed in all HCs, the top 130 (> 10.0 RPKM in all HCs) are shown in Figure 6A. An enrichment analysis, using the ShinyGO TF Targets database showed significant enrichment (FDR value <0.05) of target genes in the ELK1, GABP, PEA3 (ETV4), SP1, and STAT1 pathways. Further comparison of similarities among the HC populations showed seven TF genes co-expressed in zHCs and IHCs (not OHCs), 17 TF genes co-expressed in zHCs and OHCs (not IHCs), and 28 TF genes co-expressed in IHCs and OHCs. For comparison, we also examined the top 20 expressed TF gene orthologs in each HC type (Fig. 6B). There was notable overlap between those TF genes expressed in IHCs and OHCs, with 11 orthologs co-expressed in the list of the top 20 TFs.

**Figure 6.**
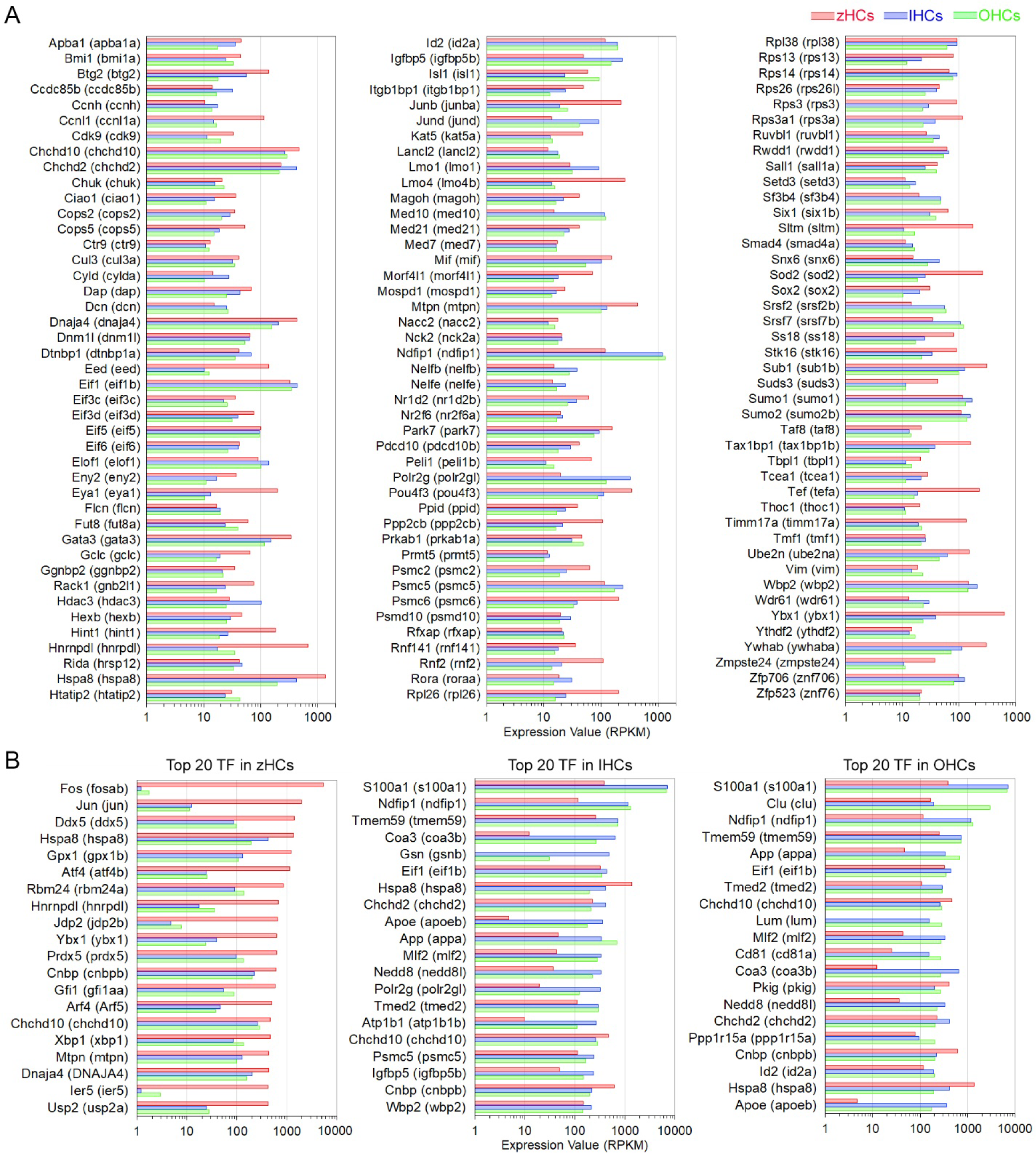
Expression of protein-coding gene orthologs categorized as transcription factors (TFs). **A**: Top 130 TF genes, classified as high-confidence orthologs, expressed in zHCs, IHCs, and OHCs (> 10.0 RPKM). **B**: The top 20 expressed TF genes in each zHCs, IHCs, and OHCs.

### Verification of gene expression in zebrafish and mouse inner ear cells

The expression of a subset of genes detected by RNA-seq (Figure 7A), selected based on unique expression patterns in zHCs, IHCs, and OHCs, was validated by RT-PCR (Figure 7B), smFISH (Figure 7C, D), and immunostaining (Figure 7E) in mouse and zebrafish tissue. As shown in Figures 7A and B, *Kcnj13*, *Ano3*, *Homer2*, *Kcnq4*, and *Chrna10* are expressed in mouse HCs. However, the corresponding orthologs have little to no measurable expression in zHCs. Positive labeling with smFISH also showed *Kcnj13* mRNA expression in OHCs, with lower expression observed in IHCs, and minimal *kcnj13* expression in zHCs (Figure 7C). Conversely, expression of *Cacnb3* in IHCs and OHCs is quite low, relative to the high-level expression of the ortholog *cacnb3b* in zHCs (Figure 7B, D). Gene orthologs uniquely expressed in zHCs include *tmc5* and *kcnq5a* (Figure 7B). Expression of genes encoding cholinergic receptors *Chrna9* and *Chrna10* in both IHCs and OHCs differs from the sole expression of *chrna9* in zHCs (Figure 7A-C). Comparatively, *slc17a8*, which encodes a vesicular glutamate transporter, is highly expressed in zHCs (Fig7D), similar to expression of the ortholog *Slc17a8* in mammalian HCs (Liu et al., 2022). Finally, we used immunostaining to examine expression of SLC7A14, a protein that is evolutionarily conserved and expressed in photoreceptors, subcortical neurons, and vertebrate HCs (Lein et al., 2007; Jin et al., 2014; Li et al., 2018). Loss of function of this gene leads to auditory neuropathy and retinitis pigmentosa in mice and humans (Giffen et al., 2022). As shown in Figure 7E, SLC7A14 is robustly and specifically expressed in adult mammalian IHCs, while the ortholog, Slc7a14, is highly expressed in zHCs. Therefore, this protein-coding gene ortholog can serve as a marker for all HCs in nonmammals and IHCs in adult mammals.

**Figure 7.**
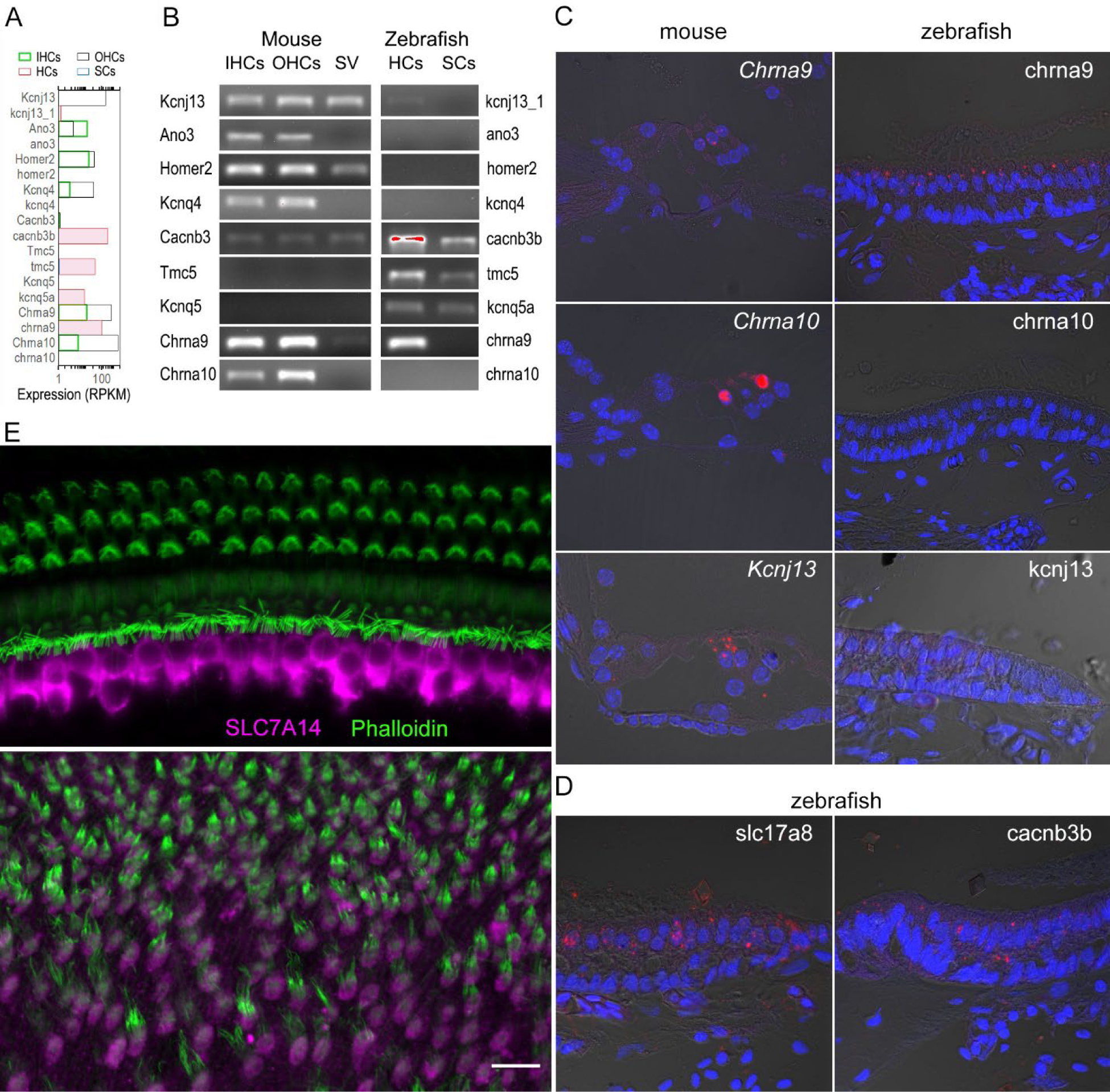
Validation of select genes expressed in zebrafish and mouse HCs. **A**: Quantified expression of select genes based on published RNA-seq data. **B**: Confirmed expression of select genes in mouse IHCs, OHCs, SV, as well as zHCs and zSCs using RT-PCR. **C**: Molecular expression of select gene orthologs visualized using smFISH in mouse (left) and zebrafish (right) inner ear tissues. (Target probe: red; DAPI: blue). **D**: Expression of *slc17a8* and *cacnb3b* in zHCs detected by smFISH. **E**: Confocal images showing expression of SLC7A14 in mouse IHCs (top panel) and Slc7a14 in zHCs (bottom panel). Scale bar (applied to all images in C to E): 10 μm.

### Electromotility and nonlinear capacitance are properties unique to mammalian OHCs

Prestin, encoded by *Slc26a5*, is the motor protein that drives electromotility of mammalian OHCs. A prestin ortholog, zprestin, was detected from the inner ear tissue of zebrafish (Albert et al., 2007). We examined *Slc26a5* expression in zebrafish and mouse HCs. As shown in Figure 4B, *Slc26a5* is highly expressed in OHCs and weakly expressed in IHCs with a ratio difference (in log2 scale) of ∼29. Weak expression of *slc26a5* is detected in zHCs. Although non-linear capacitance (NLC), an electrical signature of electromotility, was detected in a cell line transfected with the zprestin ortholog (Albert et al., 2007; Tan et al., 2011), motility and NLC have never been measured from zHCs. We measured motility and NLC simultaneously from zHCs, with similar measurements from OHCs as a positive control and IHCs as a negative control. Zebrafish HCs in utricle, saccule, and lagena have differing cell morphologies, with representative images shown in Figure 8B. Moreover, even within the same end organ, HCs located in striolar and extrastriolar regions differ in morphology and electrophysiology (Olt et al., 2014). We measured motility and NLC from 15 zHCs with different morphologies to capture regional diversity. While OHCs show robust voltage-dependent motility and large bell-shaped NLC, the zHCs do not show any signs of electromotility (Figure 8C). Capacitance response was also flat with no apparent voltage-dependence within the voltage range measured, which is somewhat different from a weak voltage-dependent response seen in *slc26a5*-transfected cell lines (Albert et al., 2007; Tan et al., 2011). Similar to zHCs, IHCs also do not show any signs of motility and NLC, despite weak expression of *Slc26a5* in IHCs.

**Figure 8.**
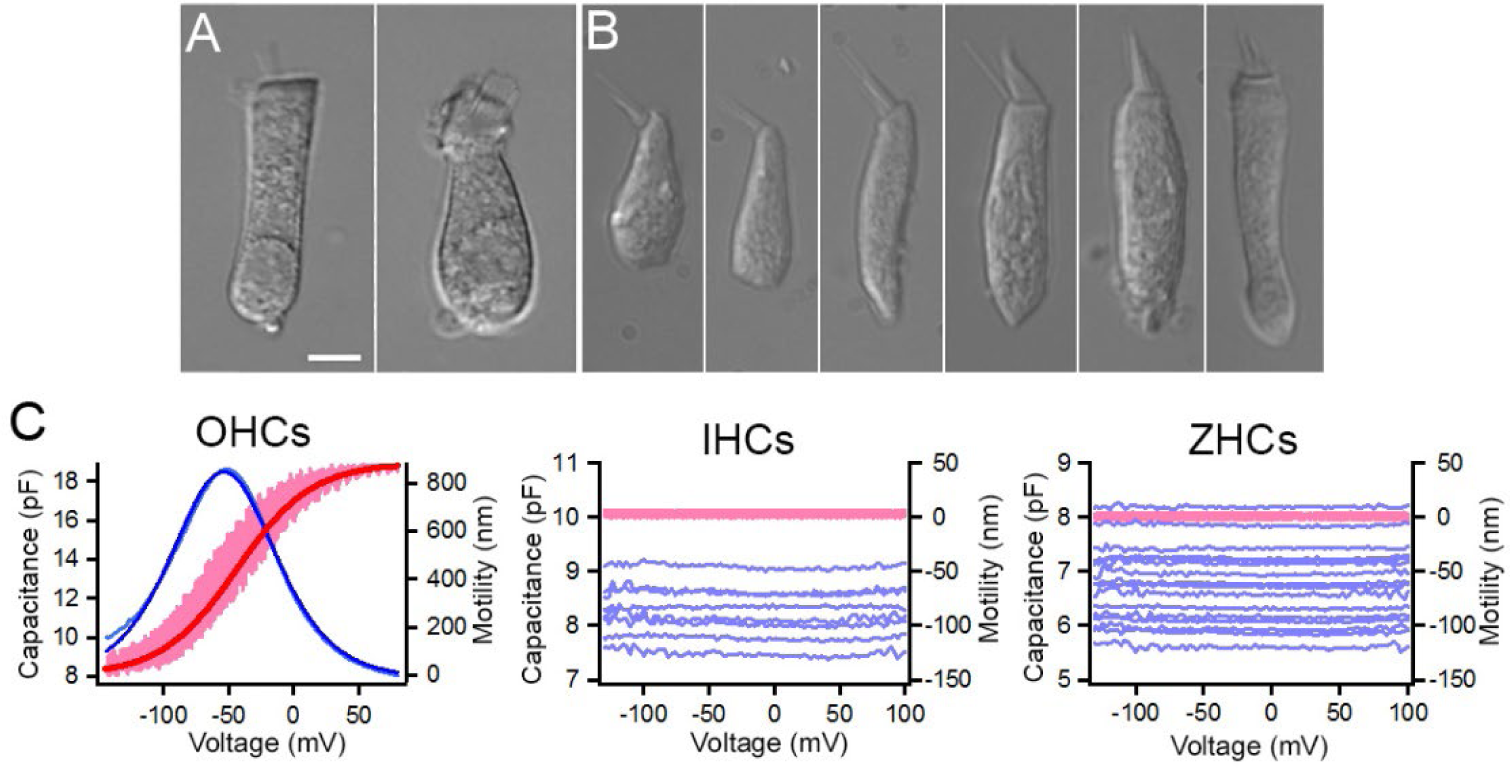
Non-linear capacitance (NLC) and motility measured from zHCs, IHCs and OHCs. Representative images of isolated **A**: mouse OHC and mouse IHC, and **B**: zHCs. **C**: NLC and motility measured from mouse OHCs (*n*=6) and IHCs (*n*=8), and zHCs (*n*=15). The pink lines represent motility, and the dark red line represents motility response fit with a two-state Boltzmann function, while light blue lines represent NLC and dark blue line represents NLC fit with derivative of a first-order Boltzmann function (Q_max_: 1135 fC, α= 27 mV^−1^, V_1/2_= −54 mV, C_lin_: 7.9 pF).

## Discussion

The challenges of depicting the molecular signatures underlying HC physiology in mammals are numerous, thus, non-mammalian model systems, including zebrafish and chicken, have become useful to explore processes such as HC development and regeneration. Numerous studies previously characterized the gene expression profiles of vertebrate inner ear HCs (Cai et al., 2015; Schrauwen et al., 2016; Barta et al., 2018; Matern et al., 2018), including some of the first studies to isolate and sequence pure populations of mammalian IHCs and OHCs (Liu et al., 2014; Li et al., 2018). More recently, single-cell RNA-seq analyses of vertebrate inner ear HC populations have exponentially increased our knowledge of the transcriptomic signatures of these specialized sensory cells. Additionally, some studies have compared gene expression homology among avian and mammalian inner ear cells, though they were not HC-type specific (Wu et al., 2023). We examined the homology of mouse cochlear IHCs and OHCs, and zebrafish inner ear HCs based on the expression of protein-coding gene orthologs among the cell populations. Thus, our study provides a novel comparative dataset to further the utility of zebrafish as a model for exploring mammalian HC function. Notably, common and uniquely expressed gene orthologs in mouse and zHCs, reveal transcriptomic signatures underlying phenotypic cellular specializations.

### Shared and unique expression of gene orthologs reveals conserved transcriptomic signatures underlying vertebrate HC specialization

To aid in understanding some of the molecular signatures common and unique to inner ear HCs, we first compared expression patterns of protein-coding gene orthologs among each cell population. While the total number of genes in zHCs is greater, due to paralogous genes, the total percentage of gene orthologs expressed in each HC population accounted for about 62 to 65 percent of the total protein-coding gene orthologs. The shared expression of genes in homologous cell populations are more likely those associated with transcriptional regulation or metabolic pathways, while there may be greater divergence in expression of genes in cellular communication (Alam et al., 2020). When examining the number of commonly expressed genes among the distinct HC populations, mammalian IHCs and OHCs shared the highest number of expressed genes. Though surprisingly, OHCs and zHCs shared more commonly expressed genes than IHCs and zHCs. Perhaps the phenotypic diversity among zHCs is captured both in the shared overlap with OHCs with their unique features including electromotility, and IHCs as the true sensory cell of the auditory system (Dallos, 1992). There were over 2,800 uniquely expressed genes in zHCs, which suggests that some of these genes encode proteins specific to this cellular phenotype or are paralogs that may have acquired a subfunction of the ancestral gene (Rohmann et al., 2009).

### Differential gene expression reveals transcriptomic signatures of non-mammalian inner ear HCs and distinctive similarities to mammalian HCs

To provide a comprehensive overview of expressed protein-coding gene orthologs in zebrafish inner ear HCs compared to zSCs, we examined uniquely expressed and upregulated genes in zHCs. Compared to the transcriptomic signatures of zSCs, the zHCs uniquely expressed several genes or gene paralogs with known contribution to HC function. Furthermore, several of the highly expressed orthologs, including *gfi1aa*, *lhfpl5a*, *myo6b*, *strc*, and *s100s* were recently described as molecular markers to delineate HC subtypes in the zebrafish inner ear (Shi et al., 2023), while others are known pan-HC genes or have been well-described as essential to HC function. For example, the high-confidence orthologs *clic5a (Clic5),* encodes an intracellular chloride channel protein that localizes to the basal region of the hair bundle and loss-of-function is associated with dysmorphic stereocilia, and vestibular and hearing dysfunction (Gagnon et al., 2006). The *cib3* ortholog, encoding calcium-binding protein, has been shown to be co-expressed with *tmc1* in the zebrafish inner ear HCs, with the ortholog *Cib3* required for mechanotransduction in cochlear HCs (Liang et al., 2021; Giese et al., 2023; Smith et al., 2023). Expression of the gene orthologs *anxa5a* and *Anxa5* is highly conserved among vertebrate HCs, despite unknown functional significance of the protein in auditory HCs (Krey et al., 2016).

### Expression of gene orthologs among zebrafish and mouse HCs reveals molecular signatures underlying sensory HC functional specialization

To further delineate the common transcriptomic signatures underlying vertebrate HC specialization, we compared expression of gene orthologues encoding functional HC proteins. For example, *Otof* (*otofa/b*), *Slc1a3* (*slc1a3a), Slc17a8* (*slc17a8*), and *Slc7a14* (*slc7a14b*) are highly expressed in zHCs and IHCs. *Slc26a5*, encoding the motor protein prestin (Zheng et al., 2000), is highly expressed in OHCs and detected at similarly low levels in IHCs and zHCs (*slc26a5*). Subsequently, we demonstrated here that zHCs do not exhibit properties of electromotility.

A great degree of conserved expression was observed among those genes orthologs encoding stereocilia-associated proteins, as over 75 percent of the genes examined were expressed in all HCs. There are few distinct differences in expression, for example *actn4* and *vill* were not detected in zHC, while both IHCs and OHCs expressed the orthologs *Actn4* and *Vill*. The absence of conserved expression in zHCs is surprising, as the actin crosslinker proteins encoded by these orthologs were also previously detected in chicken HC stereocilia (Shin et al., 2013; Avenarius et al., 2017). Conversely, other genes were highly expressed only in zHCs, such as *capza1a and 1b*, while a related ortholog *Capzb* (*capzb*) was highly expressed in all HCs. CAPZ, a heterodimeric capping protein regulating stereocilia length and width in chick and mouse cochlear and utricle HCs, can consist of CAPZA or CAPZB subunits (Avenarius et al., 2017). This warrants further exploration of the composition of these Capz subunits in zHCs and whether their interactions with other stereocilia-associated proteins such as Grxcr1, Eps8l2, Eps8/Eps8b, and Twf2a/Twf2b are conserved.

In zebrafish and mouse HCs, there was also a high degree of conserved gene expression encoding the mechanotransduction components. TMC1 and TMC2 are subunits of a transmembrane mechanosensitive ion channel that mediates mechanotransduction in vertebrate HCs. These TMC proteins, together with other proteins such as calcium and integrin binding 2 and 3 (CIB2/CIB3), accessory protein LHFPL5, and the tip-link PCDH15/CDH23 complex, are essential for mechanosensory transduction of HCs (Holt et al., 2021; Liang et al., 2021). Zebrafish HCs expressed *ush1c*, *tmie, pcdh15a/b* and *lhfpl5a*, but not the paralog *lhfpl5b*, while mouse HCs expressed the orthologs *Ush1c*, *Tmie*, *Pcdh15*, and *Lhfpl5*, respectively. Notably, zHCs expressed *tmc1*, *tmc2a* and *tmc2b*, while IHCs and OHCs expressed only *Tmc1*. *Cib2* was highly expressed in IHCs and OHCs, while both *cib2* and *cib3* were expressed in zHCs. Patterned co-expression of *tmc* subtypes with *cib2* and *cib3* among extrastriolar and striolar zHCs was recently described (Smith et al., 2023). Describing the identities and functional interactions among the components of the mechanotransduction channel in stereocilia continues to be an area of research interest and, with their underlying similarities, zebrafish can serve as an ideal model organism.

Differential expression of ion channels among vertebrate HC populations has also been an ongoing area of inquiry, often further complicated by the many splice variants encoded by ion channel genes and the subunit composition of functional ion channels. We found that zHCs, compared to their mammalian counterparts, have a more diversified pattern of expression of ion channel genes, many of which are paralogous genes. It is unclear whether the expressed genes are redundant or if the protein encoded by a gene paralog became functional during evolution, though it should not be assumed that it retained the same molecular function (Koonin, 2005). For example, zHCs express *kcnn1a* and *kcnn1b*, along with the paralog *kcnn2*. The ortholog *Kcnn1* has low expression in IHCs and OHCs, and *Kcnn2* is highly expressed in OHCs. *Kcnn2* encodes a calcium-activated potassium channel (SK2) which, along with α9α10-containing ionotropic receptors, contributes to post-synaptic organization and cholinergic inhibition of efferent synaptic cisterns of mouse OHCs (Fuchs et al., 2014). Related expression patterns among zHCs include the differential expression of paralogous genes encoding the cholinergic receptor nicotinic (nAChR) α9 and α10 subunits. Both *Chrna9* and *Chrna10* are expressed in IHCs and OHCs, while only *chrna9* is expressed in zHCs. Consistent with mammalian expression, *Chrna9* and *Chrna10* are expressed by both short and tall HCs in the chicken cochlea (Janesick et al., 2021). In birds and mammals, the α9 nAChRs are functionally coupled to SK2 potassium channels in HCs, and SK1 and SK2 are often co-expressed in the brain and can form heteromeric channels (Dulon et al., 1998; Matthews et al., 2005; Carpaneto Freixas et al., 2021). The functional properties of the co-expressed paralogous SK channels and nAChRs in zHCs warrants further investigation, especially considering the greater sequence divergence among the HC α9α10 nAChRs across species (Marcovich et al., 2020). However, *Chrna10* appears to be an adaptation of both avians and mammals.

While many of these protein-coding gene orthologs may have conserved function in vertebrate HCs, there are other unique properties in non-mammalian HCs, such as electrical resonance, that may be attributed to some of the uniquely expressed ion channel genes. This process relies on electrical resonance properties that arise from the interplay of voltage-gated calcium channels and BK channels, together with membrane properties. For example, *cacnb3b* was strongly expressed in zHCs, compared to mild expression of *Cacnb3* in mouse HCs. Similarly, *kcnma1a*, encoding a large-conductance calcium activated potassium (BK) channel, was strongly expressed in zHCs, like the orthologous gene *kcnma1* (*cslo1*) detected in both tall and short chicken HCs (Janesick et al., 2021). The high-level expression of these genes in zHCs, and not in mouse HCs, may contribute to the differential physiological properties underlying frequency tuning in non-mammalian vertebrate HCs. Differential expression of BK channel isoforms along chicken basilar papilla HCs mediates tonotopic frequency sensitivity, and knockdown of *kcnma1a* encoded BK channel expression increased auditory thresholds in larval zebrafish (Rosenblatt et al., 1997; Rohmann et al., 2014). Whereas in mammals, BK knockout mice showed normal cochlear function and increased resistance to noise-induced hearing loss (Pyott et al., 2007). The unique contributions of the ion channels encoded by *cacnb3b* and its interplay with expressed isoforms encoded by *kcnma1a* in mediating electrical resonance of zHCs warrants further investigation. Furthermore, exploring the physiologic contributions of ion channels encoded by other uniquely expressed genes in zHCs may further our understanding of the divergent specializations in non-mammalian HCs.

### Comparative transcriptomic analysis reinforces the utility of the zebrafish model for exploring hearing loss phenotypes and mechanisms of HC regeneration

While this study included only a subset of genes known to be associated to hearing loss phenotypes, those identified as one-to-one orthologs are more likely to have conserved functions. Considering this homology, characterization of induced gain or loss of function variants in zebrafish may be achieved more efficiently than mammalian models, allowing the molecular dysfunctions underlying hearing loss phenotypes to be further elucidated. For example, the lysosomal SLC transporter encoded by the orthologs *Slc7a14* and *slc7a14b* is highly expressed IHCs and zHCs, respectively, and loss of function causes progressive auditory neuropathy and retinopathy (Giffen et al., 2022). A recent study suggested that the SLC7A14 transporter had amino acid-sensing activity and mediated lysosomal uptake of GABA and subsequent inhibition of mTORC2 in hepatocytes (Jiang et al., 2023). With its distinctive expression in HCs the molecular function of SLC7A14 can be examined using a zebrafish model.

Finally, analysis of the orthologous TFs regulating gene expression in mature mouse and zebrafish inner ear HCs further elucidated key molecular similarities and differences in these cell populations. While several of the commonly expressed genes are categorized as basal cell TFs, some are well characterized HC TF genes including *Eya1*, *Gata3*, *Pou4f3*, *Nr2f6*, *Sox2* and others (Elliott et al., 2021). Among the top expressed TF genes in HCs, were several regulated by SP1 which is known to induce transcriptional responses to physiologic or pathologic stimuli. Upregulation of SP1 and subsequent up or downregulation of target genes protected cochlear HCs from oxidative damage (Li et al., 2023). STAT1 target genes, which regulate processes such as cell cycle arrest and immune response, were also enriched among all HCs. Decreased expression of STAT1 target genes reduced cisplatin-mediated apoptosis and aminoglycoside-induced inflammation in the mammalian cochlea (Levano and Bodmer, 2015; Kaur et al., 2016). More recent findings showed the therapeutic effects of compounds that ameliorate ototoxic drug-induced damage to the mouse cochlea and zebrafish lateral line HCs via regulation of STAT1 and STAT3 signaling (Yin et al., 2023; Yang et al., 2024). *Stat3 (stat3)* had lower expression in all HCs, while *stat1a* was more highly expressed in zHCs and zSCs. Given the interplay between STAT1 and STAT3 target genes and pathways, further study to elaborate why these STAT1 target genes are upregulated in all HCs, as well as the regulatory activity of Stat1 in zHCs warrants further exploration. Another group of enriched TFs included ETV target genes; ETV TFs ETV4 and ETV5 are regulated downstream of mesenchymal FGF signaling to regulate cochlear length during development (Ebeid and Huh, 2020). Both zHCs and zSCs express *etv4 (pea3)*; upregulation of FGF signaling via Wnt activation significantly upregulated *etv4*, and subsequently induced cell proliferation in lateral line neuromasts (Tang et al., 2019). Interestingly, we found that *Etv5* (*etv5a/b*) was expressed in adult mouse and zHCs, however, the role of the ETV TFs in maintaining the adult sensory epithelia is unclear.

Continued exploration of the transcriptional landscapes regulating the mature HC phenotype will further the pursuit to regenerate adult mammalian HCs. Unlike their adult mammalian counterparts, mature zebrafish and chicken HCs retain the ability to regenerate by conversion from SCs to restore function after damage (Choi et al., 2024). Regeneration has been widely studied in zebrafish lateral line HCs and some of the molecular mechanisms regulating this process have been successfully utilized to induce regeneration of neonate mammalian HCs (Ye et al., 2020; Li et al., 2022). Although overexpression of a cocktail of TFs can convert SCs to HCs in adult mammals, the challenge to fully convert these to mature, functional HCs remains (Lee et al., 2020). Additional analyses, which correlate gene expression profiles during zHC regeneration, to both developing and mature mammalian HC gene expression profiles, will provide further clues to HC regeneration in adult mammals (Baek et al., 2022). Thus, the presented dataset, including expressed TF gene orthologs in mature inner ear HCs, may serve as a transcriptomic guide for regenerative studies.

### Limitations

We recognize that describing functional homology among vertebrate HCs based on transcriptomic analyses is not without its caveats. Firstly, gene expression is not equivalent to protein expression, thus, further proteomic analysis of these and other vertebrate HC populations will further support shared and unique molecular functions. The original RNA-seq analysis of isolated zHCs were pooled from the utricle, saccule, and lagena, thus distinct gene expression patterns among the zHC subtypes are not exhibited in this study. Further analysis of orthologous gene expression defining molecular subtypes is increasingly possible as single-cell RNA-seq technologies continue to improve our abilities to isolate and sequence mature vertebrate inner ear HCs.

## Conclusion

While we focused most of this analysis on genes with known HC functions, this is by no means a comprehensive analysis of protein-coding orthologs that may contribute to ion transport, mechanotransduction, synaptic signaling, or maintenance and regeneration of the sensory HCs. Importantly, these data provide a resource for further exploring the molecular signatures underlying HC-specific functions in zebrafish and mouse. Additionally, this work creates a foundation for future comparisons of protein-coding gene ortholog expression in other species to further elucidate the conserved molecular signatures among vertebrate inner ear HCs.

## Supporting information

PCR primer sequences

## Author Contributions

KG: Conceptualization, Data curation, Formal analysis, Investigation, Methodology, Validation, Visualization, Writing - original draft and review & editing. HL: Investigation, Methodology, Visualization, Writing - review & editing. KY: Investigation, Methodology, Visualization, Writing - review & editing. YL: Investigation, Methodology, Writing - review & editing. LC: Investigation, Methodology, Writing - review & editing. KK: Methodology, Resources, Writing - review & editing. MZ: Methodology, Resources, Writing - review & editing. DH: Conceptualization, Methodology, Funding acquisition, Supervision, Project administration, Validation, Visualization, Writing - original draft and review & editing.

## Supplementary Material

Supp Data Sheet 1: Downloadable Excel file with the quantified cell-type specific protein-coding gene ortholog expression data.

Appendix 1: RT-PCR Primers

## Data Availability Statement

The datasets analyzed for this study NCBI SRP113243 and NCBI SRP133880 can be found in the NCBI GEO repository https://www.ncbi.nlm.nih.gov/geo/.

