## Supplementary material for "Molecular Specializations Underlying Phenotypic Differences in Inner Ear Hair Cells of Zebrafish and Mice": PCR primer sequences

**Appendix 1: Primer Sequences**

| **Mouse** | Forward | Reverse |
| --- | --- | --- |
| *Kcnj13* | TGACTGTCCAAGTGCAATCG | CACCTGTGATAAAAGCCTCTAGC |
| *Ano3* | TACTTCGGAGGAGAGTAGAAAGG | GGATGCAGGCTTATCTTTGGT |
| *Homer2* | ACTTTCACCAAAACGTCACAGA | GGAGAATCCCAAACCGAACAC |
| *Kcnq4* | TCAGGACTGTAGTGGATGTAGCGAC | CTGTGGATGTGTTTCCTGATGCTG |
| *Cacnb3* | GGTTCAGCCGACTCCTACAC | GAGAACTTCAAGACTGCCTGCATCC |
| *Tmc5* | GGACAGAAACTAATTGCATCCCT | GGTCCTGCGGTAAGTCAGTG |
| *Kcnq5* | CCCAAGTGGAGAGAATCCAA | TTGACGTGAGGCAAACTGAG |
| *Chrna9* | CGTGTGATCTCCACCAGTGT | TCCTTCATCCCTTTATCCTTGA |
| Chrna10 | AGCCCTTCTGCATCACGTAG | AAAGCGGTCCATTACTCTGG |
| **Zebrafish** | Forward | Reverse |
| *kcnj13* | CTGGCTGCTGTTTGCTGTATTATGG | TTGGCTACAAATGCACCAGTGATG |
| *ano3* | TCGTTCATCATGTGCTTCAGCGGAC | CAGCGTGCCCATCTCTCGTACAG |
| *homer2* | AACGGCACCGACGACGAGAAGATC | TGGTTGTTCTCCTCCAGGCTCC |
| *kcnq4* | TCAGGACTGTAGTGGATGTAGCGAC | CTGTGGATGTGTTTCCTGATGCTG |
| *cacnb3b* | TCGAGAGGGCCAAGACCAAACCTG | TTTCCTCTGAGCTGCTTTCTGTTCC |
| *tmc5* | TGCGAGAGCAGATTGTAAATGGAGG | TCCAGTTCGGGTTGTAGTATGTCG |
| *kcnq5a* | AAACTCCAATCTCACGGCGG | TCACAGTTCCTGACAAGTGC |
| *chrna9* | GGAGTCGGCTACCTTCACGATAATG | CCTGTCCATCACCTTGGCAACCTTC |
| *chrna10* | GCCTCAGGACTGCAACTGCAACATG | TTTCCAGTCTCAGCCGCTCCATCG |
